# Occurrence and predictive utility of isochronal, equiproportional, and other types of development among arthropods

**DOI:** 10.1101/379164

**Authors:** Brady K. Quinn

## Abstract

In isochronal (ICD) and equiproportional development (EPD), the proportion of total immature (egg, larval, and/or juvenile) development spent in each stage (developmental proportion) does not vary among stages or temperatures, respectively. ICD and EPD have mainly been reported in copepods, and whether they occur in other arthropods is not known. If they did, then rearing studies could be simplified because the durations of later developmental stages could be predicted based on those of earlier ones. The goal of this study was to test whether different taxa have ICD, EPD, or an alternative development type in which stage-specific proportions depend on temperature, termed ‘variable proportional’ development (VPD), and also how well each development type allowed later-stage durations to be predicted from earlier ones. Data for 71 arthropods (arachnids, copepod and decapod crustaceans, and insects) were tested, and most (85.9 %) species were concluded to have VPD, meaning that ICD and EPD do not occur generally. However, EPD predicted later-stage durations comparably well to VPD (within 19-23 %), and thus may still be useful. Interestingly, some species showed a ‘mixed’ form of development, where some stages’ developmental proportions varied with temperature while those of others did not, which should be further investigated.

**Highlights:**

- Whether arthropod development is generally isochronal or equiproportional was tested
- Developmental proportions of most species’ stages varied with temperature
- Many species had ‘mixed’ development between variable and equiproportional types
- The general occurrence of isochronal and equiproportional development was rejected
- Equiproportional development did make reasonable predictions of stage durations

## 1. Introduction

### 1.1. Importance of studying immature arthropod development

The immature phases of arthropod life cycles (i.e. eggs, larvae, and juveniles) are extremely important because they mediate recruitment to all later life stages (i.e. sexually mature adults). Survival through early life stages, growth rates of individuals to reproductive maturity or legal size for fisheries, inter-population connectivity, and seasonal patterns of abundance are all impacted by the length of time required to complete the immature phases (Huntley and López, 1992; Miller et al., 1998; Anger, 2001; Easterbrook et al., 2003; Reitzel et al., 2004; Pineda and Reyns, 2018). Development time of arthropods is strongly impacted by environmental temperature, with warmer conditions generally resulting in shorter development times within a species’ tolerance limits (Nietschke et al., 2007; Shi et al., 2012; Rebaudo and Rabhi, 2018). There is therefore much interest and need to quantify the relationships between temperature and development time of immature arthropod life stages. Once such relationships have been determined for a given species, they can then be used to derive functions that are incorporated into models of larval dispersal, life history, production, etc. so that predictions of species’ ecology and population dynamics can be made and tested against nature (Miller et al., 1998; Reitzel et al., 2004; Quinn et al., 2013). These efforts can inform fields such as fisheries ecology, food web modeling, predicting the spread of invasive species, pest management for agriculture, food science, epidemiology, forensic science, and so on (Huntley and López, 1992; Easterbrook et al., 2003; Lefebvre and Pasquerault, 2004; de Rivera et al., 2007; Yamamoto et al., 2014; Quinn, 2017).

The usual approach used to quantify temperature-dependent development relations for immature arthropods is to rear eggs, larvae, and/or juveniles at several different constant temperatures in a laboratory setting, observe the time required to complete the immature phase(s) at each temperature, and then derive a generalized equation(s) based on these observations to predict development time at any temperature in nature (e.g., Anger, 1983, 1984, 1991, 2001; MacKenzie, 1988; Easterbrook et al., 2003; Leandro et al., 2006a, b; Ouellet and Chabot, 2005; Quinn et al., 2013). While simple in principle, many issues complicate this approach. One particularly large challenge is the fact that most arthropod life history phases consist of multiple immature stages, each separated by a moult and differing in morphology, physiology, and behaviour (Anger, 2001; Li, 2002; Jacas et al., 2008; Jafari et al., 2012; Martin et al., 2014). For the majority of species, the duration of each stage differs from that of the others, with later stages tending to be longer than earlier ones (Corkett, 1984; MacKenzie, 1988), but not always. In species with this type of development, the duration of each stage must usually be observed at each temperature to model development accurately. This is problematic because it requires costly maintenance of culture conditions and checking on experimental animals for extended time periods. Importantly, this approach also often results in uncertain estimates of later stages’ durations due to small sample sizes following larval mortality in laboratory conditions (e.g., Ford et al., 1979; Klein Breteler 1980; Quinn et al., 2013; Quinn, 2017). An approach that might allow for reduced rearing costs and efforts and provide larger sample sizes for quantification of late-stage durations would thus be useful.

### 1.2. Introduction to isochronal and equiproportional development

Two developmental concepts developed through observations on larval development of copepod crustaceans seem to provide such potential shortcuts. These are the concepts of isochronal development (ICD) and equiproportional development (EPD) (Hart 1990, 1998; Petersen, 2001). In a species with ICD, the durations of all developmental stages, and thus the proportion of total development spent in each stage, are identical (i.e. the ratios the of durations of all different stages or phases to one another is 1.0) (Miller et al., 1977; Corkett, 1984; Fig. 1A). In a species with EPD, the proportion of total immature development time spent in each larval phase and stage (i.e. the ratios of the durations of different stages or phases to one another) is constant within a species (Corkett and McLaren, 1970; Corkett, 1984; Fig. 1B), and perhaps even within larger taxonomic divisions (e.g., interspecific equiproportionality (ISE) within a genus, or intergeneric equiproportionality (IGE) within a family, *sensu*Hart, 1990). Importantly, within a species with ICD or EPD the proportion of total development spent in each stage or phase is independent of rearing temperature, and thus these proportions will be constant even if different temperatures lengthen or shorten the duration of each stage, phase, and total developmental duration (Corkett, 1984; Hart, 1990, 1998); this concept is illustrated in Fig. 1A, B, E, and F. ICD and EPD have historically been viewed as species-specific traits (Hart, 1994; Kiørboe and Sabatini, 1994; Petersen, 2001), such that some species have one of these types of development, while others have neither (Peterson and Painting, 1990; Carlotti and Nival, 1991) meaning that proportions of total development time varies among stages or phases and temperatures (Hart, 1998; Fig. 1C, D, G, H).

**Figure 1.**
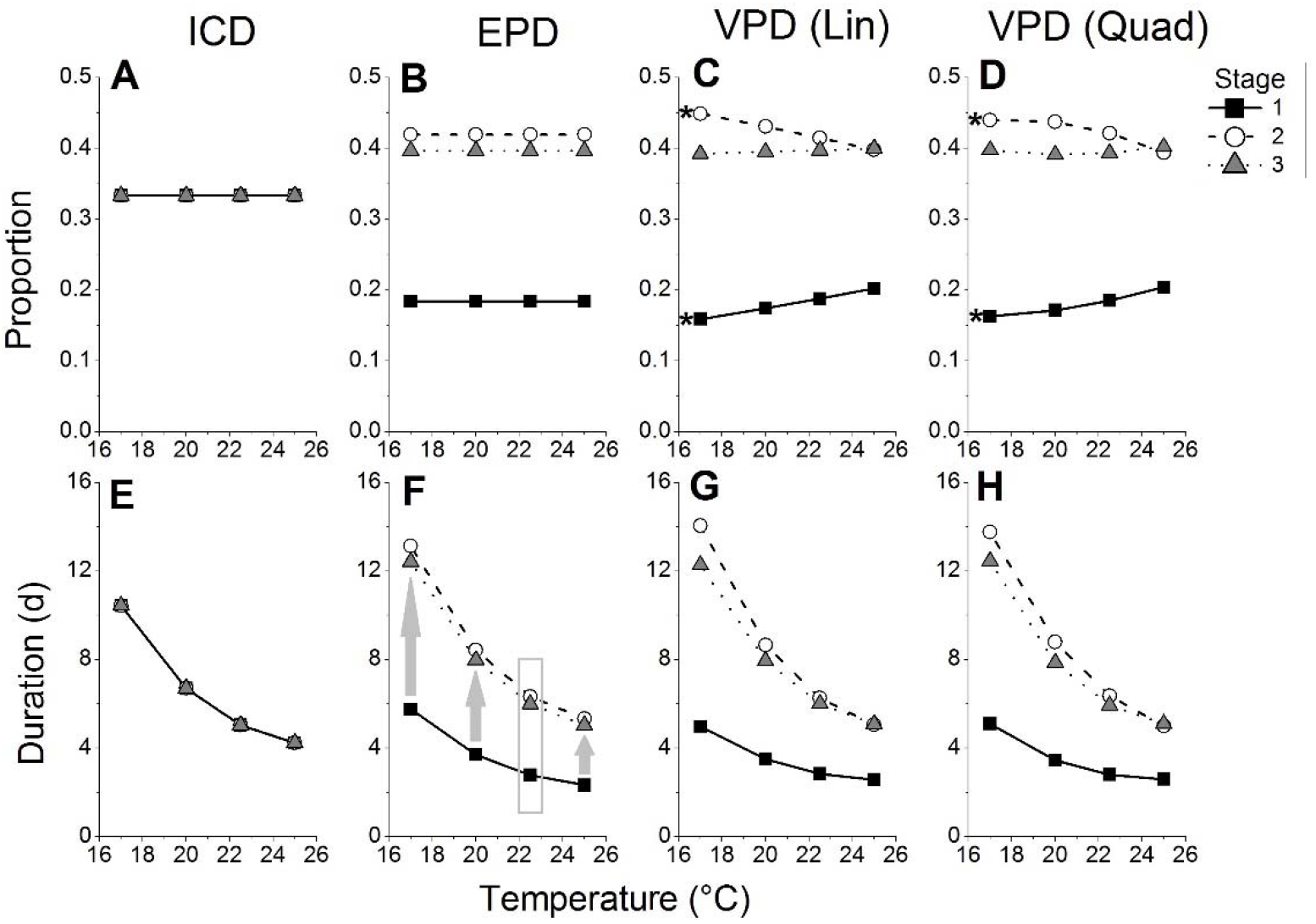
Examples of isochronal (ICD), equiproportional (EPD), and linear (Lin) and quadratic (Quad) forms of ‘variable proportional’ development (VPD) of a hypothetical arthropod with three immature stages (signified by different symbols and lines). Plots show the proportion of total development (A-D) and development time (in days (d), E-H) spent in each stage at each of four temperatures (in °C). In ICD (A, E) values for all stages are identical and overlap, in EPD (B, F) values are the same across temperatures but differ among stages, and in VPD (C, D, G, H) values differ among temperatures and stages. In (F), the use of EPD to predict the durations of later stages (2 and 3) at temperatures of 17, 20, and 25°C based on observed durations of stage 1 at these temperatures and observed proportions of development spent in all three stages at 22.5°C (gray box) is illustrated. In (C) and (D), ‘mixed’ development is illustrated, in which proportions spent in stages 1 and 2 are affected by temperature (the ‘*’ indicates a significant effect of temperature) but those in stage 3 are not.

Initial studies rearing all immature stages of a species at multiple temperatures would be needed to confirm whether a given species of interest has ICD or EPD. Once these had been done, however, subsequent studies attempting to expand on different aspects of its temperature-dependent development [e.g., rearing larvae from different source populations to test for local adaptation (Leandro et al., 2006a; de Rivera et al., 2007; Quinn et al., 2013), rearing at more extreme temperatures to assess thermal limits (Pakyari et al., 2011; Shi et al., 2012)] should be able to use these estimates and forgo the expensive rearing of later stages. In the extreme case, a species with ICD need only be reared through one immature stage at one temperature for the durations of all other stages at all other temperatures to be estimated (Fig. 1A, E), while for a species with EPD rearing all stages at one temperature could in principle provide enough information to predict the durations of all stages at other temperatures (Corkett and McLaren, 1970; Corkett, 1984; Hart, 1990; Fig. 1B, F). In practice, because low food quality [a frequent issue in larval culture (Leandro et al., 2016b)] or other stressors [e.g., low pH (Keppel et al., 2012; Quinn, 2016), extreme temperatures (Quinn, 2016, 2017)] can lead to altered stage-specific developmental proportions for a species with ICD or EPD (Hart, 1998), it would likely be safest to rear all stages at one temperature and at least one stage at several other temperatures, as recommended by Corkett (1984) (see also Fig. 1F). However, even this approach can minimize rearing costs and efforts. This could further allow the durations of later stages for individuals that die during rearing in earlier stages to be estimated, giving a larger sample size for the quantification of later-stage durations and capturing more of the natural variability therein. There are, however, important limits or uncertainties to using ICD and/or ICD in this way, specifically that: (1) it is not known whether non-copepod crustaceans or other arthropods have these kinds of development; (2) there is no unequivocal, agreed-upon technique with which to actually *test* for the occurrence of ICD or EPD versus alternative possible development modes (e.g., in which stage proportions vary with temperature); and (3) no previous study has quantified the predictive utility of using ICD or EPD proportions to forecast later-stage durations based on durations of early stages. The present study sought to investigate and remedy these shortcomings.

### 1.3. History of the study of ICD and EPD

The concepts of ICD and EPD were initially described and named by Corkett (1984) based on observations of ICD in the copepods in the genus *Acartia* by Miller et al. (1977) and EPD in the copepods *Pseudocalanus minutus* (KRØYER, 1845), *Eurytemora affinis* (POPPE, 1880) (as *E. hirundoides*), and *Temora longicornis* (MÜLLER O.F., 1785) by Corkett and McLaren (1970). Since then, these concepts (especially EPD) have been used and developed in numerous subsequent studies on copepods. Various studies of copepod species have claimed to demonstrate ICD and EPD (Hart, 1990; Petersen, 2001; Supplementary File 1, Table S1). Aside from general rearing studies, EPD studies have also investigated numerous topics, including: taxonomic patterns in the occurrence of ICD and EPD (Hart, 1990); the implications of developmental proportions to phenology-mediated interspecific interactions among sympatric copepods species (Hart, 1994); differences between egg-carrying and free-spawning species (Kiørboe and Sabatini, 1994); dependence of developmental proportions on food quality but independence from food quantity (Hart, 1998); differences between marine and freshwater copepod studies (Petersen, 2001); and impacts of toxicants affecting the moulting cycle on EPD (e.g., copper and chromium:Gutiérrez et al, 2010a), but not if they did not impact moulting (e.g., lindane: Brown et al., 2003; fish kairomones: Gutiérrez et al., 2010b).

Based on the above, it is apparent that research using the concepts of ICD and EPD has been copepod-dominated; indeed, all 21 relevant studies found in a 1 December 2017 Web of Science (Clarivate Analytics, 2018) search for ‘equiproportional development’, and all relevant studies within the first ten pages of a Google Scholar (Google, 2018) search for the same term on the same date, were carried out on copepods (Supplementary File 1, Table S1). However, there is no particular reason that these types of immature development may not also occur among other crustacean taxa (e.g., Miller, 2008), or perhaps even in non-crustacean arthropods (i.e. insects and arachnids). A handful of studies on non-copepod arthropods have applied these concepts (in-name or not) to their species’ data. Two studies of American lobster (*Homarus americanus* H. MILNE EDWARDS, 1837; Crustacea: Decapoda: Astacidea) larval development concluded that this species has EPD at temperatures from 10-22°C (MacKenzie, 1988; Quinn et al., 2013).

There is preliminary evidence suggesting that the larval development of other decapod crustaceans, such as snow crab (*Chionecetes opilio* (O. FABRICIUS, 1788); Decapoda: Anomura) and northern shrimp (*Pandalus borealis* KRØYER, 1838; Decapoda: Caridea), is equiproportional (P. Ouellet and B. Sainte-Marie, Fisheries and Oceans Canada, pers. comm.). Miller (2008) even speculated that development of the Aesop shrimp (*Pandalus montagui* LEACH, 1814; Decapoda: Caridea) might be isochronal at 12-15°C. One study of a non-crustacean arthropod that considered developmental proportions at different temperatures was done by Lefebvre and Pasquerault (2004) on the dump fly *Ophyra aenescens* (WIEDEMANN, 1830) (Insecta: Diptera). That study found that this species spent different percentages of its total immature development in each stage at different temperatures (17-30°C), and thus did not have ICD or EPD. Other studies of non-copepods or non-crustaceans considering developmental proportions spent in different stages may exist, but are difficult to find because the ICD and EPD terminology is largely unknown outside of the copepod literature. Given the potential reductions in study effort and costs for rearing a species with ICD or EPD discussed above, however, it would be worthwhile to investigate whether and which other arthropod taxa show these kinds of development.

### 1.4. Uncertainties regarding ICD and EPD: possible alternatives and predictive ability

Before various arthropod species can be investigated to determine whether or not they have ICD or EPD, an objective test is needed that can confirm or reject either of these types of development, and/or provide evidence that they described development data for a given species better than an alternative type of development (e.g., in which proportions vary among stages and temperatures; Petersen and Painting, 1990; Fig. 1C, D, G, H). Most of the foundational studies on copepods that considered ICD and EPD did so in a very informal manner, for example by calculating copepodite:nauplius or egg:nauplius phase duration ratios at different temperatures and subjectively deciding whether these were similar across temperatures (Corkett and McLaren, 1970; Corkett, 1984) or even just calculating ratios across temperatures (e.g., Hart, 1990). Such methods are not appropriate to make unequivocal conclusions regarding the type of development a given species has (e.g., Peterson and Painting, 1990; Carlotti and Nival, 1991). Studies by MacKenzie (1988) and Quinn et al. (2013) attempted to ‘test’ for EPD using formal statistical techniques, accounting for inter-individual variability and testing across temperatures. MacKenzie (1988) compared proportions using a two-way analysis of variance (ANOVA) with rearing temperature (5 levels) and larval stage (4 levels) as interacting factors. Quinn et al. (2013) performed separate one-way ANOVAs for each larval stage, with temperature as a factor with 4 levels. Both of these approaches treat temperature, a continuous variable, as a categorical factor, which is technically incorrect. Quinn et al. (2013)’s approach, or a similar alternative approach performing separate regressions or correlations of temperature vs. proportion for each stage, does not allow for comparisons among larval stages and thus cannot test for ICD vs. EPD. An analysis of covariance (ANCOVA, either linear or nonlinear), with the continuous variable temperature as ‘covariate’ and larval stage as a categorical factor may be a better general approach, although the typical interpretation of an ANCOVA focuses on the categorical factor and ignores functional relationships between the covariate or covariate*factor interaction and dependent variable. An ANCOVA approach is thus not ideal since functional quantitative relationships between temperature and developmental characteristics of arthropods are often quite important (e.g., Shi et al. 2012; Quinn, 2017; Shi et al. 2017a, b, c; Rebaudo and Rabhi, 2018).

An optimal approach would be the use of multiple generalized linear models, each designed to represent a hypothesis of development type (ICD, EPD, other), with an information-theoretic approach used to compare models and select the ‘best’ one for a given species dataset (Anderson 2008; Quinn 2017). This latter approach was adopted in the present study, as detailed in section 2 (Materials and methods). An important component of such a method is the development of an alternative model, in which developmental proportions vary with temperature, against which to compare ICD and EPD. Within the present study, this form of development was termed ‘variable proportional development’ and abbreviated ‘VPD’, and models representing it were developed based on previous research on arthropod development. A number of studies have investigated the ‘rate isomorphy’ hypothesis, which proposes that relative durations of particular developmental stages are constant across temperatures, and/or that lower developmental thresholds are constant among development stages, on various ectotherm taxa, including insects, mites, and bamboos (e.g., Jarošík, 2002, 2004; Shi et al., 2010; Sandhu et al., 2011; Kuang et al., 2012; Lin et al., 2018). In these studies, development rates were proposed to increase linearly with temperature across a range of intermediate temperatures, but could potentially deviate from linearity at more extreme (low and high) temperatures approaching a species’ or stage’s thermal limits. When such studies compared lower developmental thresholds across stages and temperatures, differences among stages but not temperatures demonstrated potential patterns comparable to EPD across many taxa, although with some potential exceptions; for example, deviations of the temperature-development relationship for a given stage(s) from linearity could indicate non-EPD (i.e. VPD) conditions. In addition, many previous studies of arthropod development have used both linear and nonlinear models to fit development time or rate data (e.g., Angilletta, 2006; Quinn 2017; Shi et al. 2017a, b, c; Rebaudo and Rabhi, 2018). Therefore, in the present study alternative VPD models to ICD and EPD were used, in which the developmental proportion for each stage could vary among stages as in EPD, but also could vary for the same stage as a linear or nonlinear (quadratic) function of rearing temperature.

If ICD, EPD, or VPD are widespread phenomena among arthropod immature development, researchers should still of course be wary of relying too heavily on estimates (e.g., Corkett and McLaren, 1970; Corkett, 1984; Hart, 1990; Fig. 1F) of larval development times based on these. Therefore, the predictive utility of such developmental characteristics to estimate actual immature durations should be assessed; specifically, can one make accurate predictions of later larval stage durations based developmental proportion and duration of earlier stages? No study has yet tested the usefulness of these concepts in this way.

In this study, I examined data from studies of temperature-dependent immature development in arthropods from various taxonomic groups (copepod and non-copepod crustaceans, arachnids, and insects). I developed a technique to test for ICD, EPD, or VPD for each species (Fig. 1A-D). The distribution of different development types and the performance of models based on different development types to describe species’ datasets across taxa were determined. The ability of each development type’s model to estimate later-stage durations based on earlier stages was also assessed. This is a unique study that applies and objectively tests for ICD, EPD, and alternative development types (VPD) of arthropods, including non-copepod species. Results have implications to rearing studies of immature arthropod stages in temperature-dependent development studies, and may also provide a first step towards obtaining information useful for further ecological and evolutionary studies.

## 2. Materials and methods

### 2.1. Sources and handling of data

To obtain arthropod developmental data for analyses, studies cited in a review by Quinn (2016) or whose data were reanalyzed in a recent meta-analysis by Quinn (2017) were examined. Studies that presented development time or rate data for multiple immature developmental stages of one or more arthropod species at different rearing temperatures, and in a way that allowed data to be extracted for analyses, were used as data sources. After excluding studies not meeting these criteria (e.g., those presenting only mean values, or only data for total development), 71 different species’ datasets from 60 different studies were obtained. While the primary objective of the present study was to test for ICD, EPD, or VPD in non-copepod arthropods, any copepod studies that presented useful data were still included to allow them to be tested using the new methods developed herein. The datasets collected came from studies published between 1963 and 2014 that examined various marine, freshwater, and terrestrial arthropods belonging to the arthropod subphyla Chelicerata (all arachnids, consisting of 9 mites and 1 spider), Crustacea (9 copepods and 19 decapods), and Hexapoda (33 insects belonging to 6 different orders) (Supplementary File 1, Table S2), and with both direct (anamorphic) and indirect (metamorphic) development (Table 1).

**Table 1.** The number of species’ datasets (in parentheses) for which data on different developmental stages and/or phases were obtained. Stages written in italicized text are true larvae, present in taxa (crustaceans and holometabolous insects) that undergo metamorphic development only; taxa without larvae (arachnids and hemimetabolous insects) undergo direct or anamorphic development through juvenile instars (e.g., nymphs). Within each column, each row represents a different developmental phase (for some taxa, there is more than type of larval and/or juvenile phase). Stage ranges listed in square brackets were reported as combined durations in the indicated number of studies.

| **Chelicerata: Arachnida** |  | **Crustacea** |  | **Hexapoda: Insecta** |  |
| --- | --- | --- | --- | --- | --- |
| **Acari** | **Aranea** | **Copepoda** | **Decapoda** | **“Hemimetabola”<sup>a</sup>** | **Holometabola** |
| **Egg (9)** | Egg (0) | Egg (7) | Egg (0) | Egg (6) | Egg (16) |
| **Juvenile: “larva”<sup>b</sup> (8), protonymph (7), tritonymph (2), deutonymph (5); [total juvenile (2)]** | Juvenile instar: I (0), II (1), III (1), IV (1), V (1), VI (1), VII (1) | *Nauplius*: I (3), II (3), III (4), IV (4), V (4), VI (4); [I-VI (5)] | *Nauplius*: [I-VI (1)] | Nymphal instar: I (8), II (8), III (6), IV (4), V (2); [egg + *instar I* (1)] | *Larval instar*: I (12), II (12), III (12), [I-III (1)], IV (9+1<sup>d</sup>), V (4), VI (2) |
|  |  | *Copepodite*: I (5), II (5), III (5), IV (5), V (5), I-V (1) | *Protozoaea*: [I-III (1)] | Prepupa (1); [instar I-V + prepupa (3)] | *Prepupa* (6); [egg + *instar I-V* + *prepupa* (1)] |
|  |  |  | *Zoea (or mysis)*: I (17), II (18), III (15), [I-III (1)], IV (8+1<sup>c</sup>), V (5), VI (2), VII (1), VIII (1), IX (1) | Pupa (2); [instar II-V + prepupa + pupa (1)] | Pupa (18) |
|  |  |  | *Decapodid* (8) | Pre-reproductive (2) |  |

Notes to Table 1: <sup>a</sup>Species belonging to subclass Paraneoptera, orders Hemiptera and Thysanoptera; <sup>b</sup>The first juvenile instar of mites is called a “larva” by convention; <sup>c</sup>For one
species, this stage was subdivided into 4 sub-stages (Zoea IVa-IVd); <sup>d</sup>For one species, this instar was divided into an active feeding sub-stage (“before turning purple”) and an inactive prepupal sub-stage (“after turning purple”); see Supplementary Files 1-3 for more information.

Developmental stages of these arthropods included any immature (pre-adult) stages, such as eggs, larval stages of crustaceans (nauplii, coepodites, zoeae, and decapodids), larval instars of insects, prepupae, and pupae of holometabolous insects, and nymphs of arachnids and non-holometabolous insects (Table 1; Supplementary File 2). Importantly, not all studies distinguished the durations of different developmental stages or instars within particular developmental phases (Table 1), but for the purposes of the present study all reported developmental periods (phases, stages, and/or instars) were equated. A phase contains multiple stages and has distinctive differences in its gross morphology from other phases (e.g., egg vs. larva vs. pupa; nauplius vs. zoea vs. decapodid) (Martin et al., 2014), while each stage is more similar to each other stage, although with smaller changes between them, and usually an instar involves an increase in size with little to no change in morphology (Martin et al., 2014). However, equating these periods was reasonable in light of historical research on ICD and EPD, in which not only stage-specific proportions but also phase length ratios (e.g., nauplius vs. copepodite) were considered (Corkett, 1984; Hart, 1990). Additionally, among insects and arachnids the term ‘instar’ has a more specific meaning than it does among crustaceans, involving morphological changes that may be comparable to shifts between crustacean larval stages (e.g., Huang et al., 2008; de Oliveira et al., 2009; Eliopoulous et al., 2010). Phases, stages, and instars are separated by a moult in most species, and thus the timing of transitions between them and their durations is likely under the control of similar processes (Anger, 2001; Martin et al., 2014; Quinn, 2016, 2017). Further, for each species, data on the full span of immature phases before sexual maturity was captured except that for some species the duration of the egg phase was not reported and durations of juvenile stages were not reported for decapod crustaceans, as is typical for this taxon (see Table 1). It is thus unlikely that any inconsistencies among phases, stages, and instars should have strongly affected the conclusions drawn in the present study.

Raw data or means ± error (standard deviation (SD), standard error (SE), etc.) of development time and sample sizes for each developmental stage (and for total development if presented) were extracted from tables or figures in published papers for each of these 71 species and used to generate datasets for analyses. Data were extracted from figures using ImageJ (Schneider et al., 2012). If total development times were not presented in published studies, these were calculated by summing the average duration of all developmental stages and using the error variability and sample size of the last developmental stage reported; this procedure was applied for ten (14.1 %) of the 71 species datasets.

For each species dataset, once development time data were extracted the duration of each developmental stage had to be converted into a proportion of total development time to test for ICD, EPD, etc. This was done by dividing the duration of each stage by the total duration of development extracted or calculated above. For several species, supernumerary (‘extra’) developmental stages occurred at some rearing temperatures, but not others (e.g., the zoea V stage of the coconut crab *Birgus latro* (LINNAEUS, 1767) occurred at temperatures ≤ 24.6°C, but not at higher temperatures; Hamasaki et al. 2009); such ‘extra’ stages were not considered in analyses in the present study (see Supplementary File 2), although they were accounted for in the calculation of total development times (thus influencing the relative developmental proportions of other ‘normal’ stages) if appropriate. Many studies also reported experimental treatments (e.g., rearing temperatures) or replicates at which some (usually later) stage durations or total developmental duration could not be reported due to poor or zero survival. Such treatments at which all developmental stages did not occur were also excluded from analyses (see Supplementary File 2 for the data included from each study). All data tallying and initial calculations were performed in Microsoft Excel 2013.

### 2.2. Testing each species’ dataset for ICD, EPD, and VPD

Each species’ dataset was put through a sequence of statistical tests designed to assess whether its development was best explained by an ICD or EPD model of development, or an alternative type of VPD. These tests were carried out in R version 3.1.1 (R Core Team, 2014) using a code by the author (provided in Supplementary File 2). Data fed into each test were replicates of the proportions of total development made up by each developmental stage at different rearing temperatures in a given study. Because the data were proportions, their square-root was first taken and then arcsine-transformed to meet assumptions of parametric tests (homogeneity of variances and normality of residuals) (Ahrens et al., 1990). Four generalized linear models (GLMs), all run assuming a Gaussian (normal) distribution of residuals (i.e. default settings), were then tested on the data using the glm function in R. (GLMs were used instead of regular linear models (lm) to allow them to be input to the cross validation module described later in this section.) Each GLM was intended to represent a different alternative mode of arthropod development (see Table 2, and also Fig. 1A-D). First, an intercept-only model was tested to represent ICD, since in a species with ICD all developmental stages would take up identical proportions of total development. Second, a model including only development stage as a categorical factor was used to represent EPD, since in a species with real EPD the proportions of different stages can vary from each other, but are not affected by extrinsic factors like temperature. Two further models of VPD were then tested that represented testable alternatives to ICD and EPD (see section 1.4; Fig. 1C, D), in which the developmental proportion for each stage could vary among stages as in EPD, but also could vary for the same stage as a function of rearing temperature. Both a linear and a nonlinear (quadratic) version of this development type were tested, and their GLMs included stage as a categorical factor and temperature as a continuous factor, as well as the interaction of stage × temperature; the quadratic form of VPD also included temperature^2^ and the interaction of stage × temperature^2^ (Table 2; see also Supplementary File 2). Although many different nonlinear function types exist that could be applied to arthropod development data (e.g., Quinn, 2017) and thus represent VPD, many of these require larger sample sizes and coverage of broader temperature ranges than those used in many of the studies examined (see Supplementary File2; also Quinn, 2017) to produce reasonable parameter estimates and fit. Therefore, in the present study a quadratic form of the nonlinear VPD model was used, as the quadratic development function is relatively simple and able to achieve fair performance on most datasets (Quinn, 2017).

**Table 2.** Generalized linear model (GLM) equations representing different modes of arthropod immature development (ICD = isochronal, EPD = equiproportional, and VPD (Lin) and VPD (Quad) = ‘variable proportional’ development following a linear or quadratic form, respectively) tested on arcsine-square-root transformed developmental proportion data in this study. In these equations Prop(S_i_, T) = the proportion of total development spent in stage ‘i’ at temperature ‘T’ (in °C), n = the total number of stages, S_2_, S_3_, …, S_n_ are logical variables with values of 1 if S_i_ is equal to the number of a given stage (S) or 0 otherwise, A-F and X-Z are fitted coefficients multiplied by the stage (S) and temperature (T) variables or their interactions, the intercept is the transformed proportion of development spent in the first developmental stage (S_1_), and a_1_ and b_1_ are fitted coefficients representing the effects of T and T^2^, respectively, on developmental proportions in general and the proportion of development spent in S_1_ specifically.

| Model | Generalized linear model equation |
| --- | --- |
| ICD | $\sin^{-1} \sqrt{Prop(S_i, T)} = \sin^{-1} \sqrt{n^{-1}} = \sin^{-1} \sqrt{Prop(S_1)}$ |
| EPD | $\sin^{-1} \sqrt{Prop(S_i, T)} = \sin^{-1} \sqrt{Prop(S_1)} + AS_2 + BS_3 + \dots + XS_n$ |
| VPD (Lin) | $\sin^{-1} \sqrt{Prop(S_i, T)} = \sin^{-1} \sqrt{Prop(S_1)} + a_1T + AS_2 + BS_2T + CS_3 + DS_3T + \dots + XS_n + YS_nT$ |
| VPD (Quad) | $\sin^{-1} \sqrt{Prop(S_i, T)} = \sin^{-1} \sqrt{Prop(S_1)} + a_1T + b_1T^2 + AS_2 + BS_2T + CS_2T^2 + DS_3 + ES_3T + FS_3T^2 + \dots + XS_n + YS_nT + ZS_nT^2$ |

Once all four GLMs had been run on a dataset, four metrics were calculated for each model to assess and compare how well each represented the data tested compared to the others. First, values of Akaike’s Information Criterion (AIC) adjusted for realistic finite sample sizes (AIC_C_), which represented the amount of useful information in each model (Anderson, 2008), were calculated using the AICc function in the AICcmodavg package in R (Mazerolle, 2016). The model with the lowest AIC_C_ value of all those tested on a given dataset is the best model for those data. Models were thus ranked from 1 (best) to 4 (worst if there were no ties) based on their AIC_C_ for each dataset. Second, once the best function for a dataset was determined according to AIC_C_ then model Δ_i_ values were calculated as the AIC_C_ of a given model minus the AIC_C_ of the best model (Anderson, 2008). The best model therefore has Δ_i_ = 0, and the Δ_i_ of other models gives an index of information lost by using them rather than the best model; in general, if Δ_i_ < 2 a non-best model still may contain some useful information, while if Δ_i_ > 14 it is highly unlikely to contain useful information (Anderson, 2008). Third, a pseudo-R^2^ value representing model fit to the data was calculated by obtaining null and residual deviance values from GLM summary output in R (Supplementary File 2), and then dividing the residual by the null deviance and subtracting this from one. It should be noted that following this approach the ICD (intercept-only) model always had pseudo-R^2^ = 0 because its residual deviance equals its null deviance; comparisons among pseudo-R^2^ values were thus made with caution. Finally, the cross-validation error (%) of each model was calculated to assess how well each model did at predicting data (Davison and Hinkley, 1997; Picard and Cook, 1984). Cross-validation error was determined for each GLM using the cv.glm function in the boot package in R (Canty and Ripley, 2016), carrying out leave-one-out cross-validation (K = 1) using the average squared error function (i.e. default parameters) (Davison and Hinkley, 1997). Overall, for each model a lower (better) ranking based on AIC_C_, lower Δ_i_, higher pseudo-R^2^, and lower cross-validation error indicated that that model performed better on the test dataset than others.

For each study dataset, the best development model (ICD, EPD, or linear or quadratic VPD) selected by AIC_C_ was concluded to represent its species’ ‘true’ mode of development (i.e. the best estimate thereof), and other parameters gave indices of how well the best and other development models represented its data. If the best model was found to be one of the VPD models, for instance, then a given species’ development could be concluded to not be equiproportional or isochronal. To further examine the consistency of development stage proportions among stages and temperatures for each dataset, GLM summary outputs from R were consulted. Whether coefficients calculated by the GLM for each factor and each level of each factor (for stages) were statistically significant (p ≤ 0.05) was examined (see Table 2 for model equations and Supplementary File 2 for R outputs). For the EPD and both VPD models, if any stage was significant this was concluded to represent a significant effect of stage on developmental proportions, and thus a rejection of ICD. For the VPD models, if temperature, temperature^2^, or the interaction term of any stage with temperature and/or temperature^2^ was significant, then proportions of one or more stage(s) significantly varied among temperatures, rejecting both ICD and EPD.

Results of the above inferences generally agreed with conclusions based on AIC_C_ rankings and applying ANOVAs to model outputs (see Supplementary File 3, Table S3), but provided additional information of interest. Specifically, for many species’ datasets some stages’ proportions significantly varied with temperature, but those of other stages did not significantly interact with temperature. Such species were concluded to have a ‘mixed’ form of development intermediate between EPD and VPD (discussed more in section 2.3). For each stage that varied among temperatures in a species with mixed development, the number of that stage in the developmental sequence as well as the proportion of development completed up to the end of each stage (the number of that stage in the developmental sequence divided by the total number of stages examined) was noted and examined later. These data were visually examined to see whether there were any patterns among taxonomic groups or in relation to the position in development of particular stages, for instance whether stages occurring earlier or later in development were more or less likely to have their proportions affected by temperature. Logistic regressions were also conducted to determine whether the likelihood of the proportion of development spent in a given stage being affected by temperature was related to either that stages’ number or the proportion of development completed up until the end of that stage.

### 2.3. Overall distribution and performance of each development type

Once the aforementioned statistical analyses and calculations had been completed for all 71 datasets, the total number of species’ datasets for which each development model type (ICD, EPD, linear VPD, and quadratic VPD) was concluded to be ‘best’ based on AIC_C_ was summed up, and the proportion of all datasets best represented by each model was calculated. The number and proportion of species’ datasets with ‘mixed’ development was also calculated. These data were also partitioned by arthropod subphylum (Chelicerata, Crustacea, and Hexapoda) to examine whether any overall differences in the distribution of development types existed among subphyla; copepods were also tallied separately from other crustaceans (see section 2.4). The overall performance of each development model (ICD, EPD, linear VPD, and quadratic VPD) and for different subphyla across datasets was also assessed by conducting two-way ANOVAs on the AIC_C_-based rankings, model Δ_i_, pseudo-R^2^, and cross-validation error values. The factors in these ANOVAs were development model (4 levels) and arthropod subphylum (3 levels), and each datum was one of the four performance metrics calculated for a given model on a given species dataset. Because pseudo-R^2^ and cross-validation error (converted from %) values were proportions, the square-roots of these data were arcsine-transformed to meet assumptions before analyses (Ahrens et al., 1990). If an overall statistically significant difference (p ≤ 0.05) was found among models or subphyla, Tukey’s Honestly Significant Difference (HSD) test was used to perform post-hoc comparisons. These analyses were carried out to assess how well each development model performed across datasets, and to determine whether there were meaningful overall differences between the best models for each dataset and the alternative models.

### 2.4. Comparison of model performance on copepods versus other taxa

As the concepts of ICD and EPD were initially developed in studies of copepods and have historically been thought to apply well to development of these crustaceans (Corkett, 1984; Hart, 1990, 1994, 1998; Petersen, 2001), it is feasible that these models might perform better on data for copepod crustaceans than they do on data for other arthropods. Therefore, an additional set of two-way ANOVAs on the four performance metrics described above (AIC_C_ rank, Δ_i_, pseudo-R^2^, and cross-validation error) was carried out, but this time the factors tested were development model (4 levels, as before) and ‘copepod’ status (2 levels), meaning whether a given species was a copepod or not. Tukey’s HSD test was used to perform post-hoc comparisons among models if there were significant differences among these.

### 2.5. Implications of development type: calculation of prediction error

One of the potential uses of ICD, EPD, and even VPD, is that if the relative proportions of total development taken up by different stages (and at different temperatures for VPD) are known, then not all developmental stages need to be raised (at least in all treatments) in future studies assessing immature development in a particular species. In the extreme case, one could for example only rear the first developmental stage under particular experimental conditions, and use knowledge of ICD, EPD, etc. and known proportions to estimate the durations of all later developmental stages based on that of the first one (e.g., Fig. 1F). The effectiveness of this method of extrapolation was examined in the present study by calculating prediction errors for each of the four development model types considered. For each dataset and each model, actual durations of the earliest examined developmental stage and proportions for all subsequent development stages based on each model were used to estimate the duration of each later stage. For example, if stage 1 duration was 10 days and based on the EPD model stage 1 takes up 0.4 of total development time while stage 2 takes up 0.6, then stage 2 duration could be estimated as 10*(0.6/0.4), or 15 days. For VPD models, variation in developmental proportions for each stage among rearing temperatures was also accounted for in these calculations.

Predicted durations were then compared to actual reported durations to estimate prediction error. To make values comparable across species with disparate developmental durations, and because no directionality was expected in terms of prediction errors, prediction error was calculated as an absolute percentage value by subtracting predicted from observed duration, dividing by observed duration, multiplying by 100 %, and taking the absolute value. For each study dataset the average prediction error of each development model was averaged across all stages and replicates, and this average value was then used in subsequent overall analyses. The magnitude of overall prediction error for each model across datasets was examined. The minimum prediction error value from which the overall distribution of prediction errors of each model was not significantly different was determined through a series of one-sample *t*-tests, in which the data were compared to test values of first 0 %, and then gradually increasing percent error values (at 1 % increments) until a p-value ≥ 0.05 was attained. Whether mean prediction errors varied among development models or arthropod subphyla across datasets was also tested with a two-way ANOVA, using Tukey’s HSD test for post-hoc comparisons among levels of any significant factor(s). Whether prediction errors of the four models differed overall between copepods and all other taxa examined was also tested with another two-way ANOVA with copepod status as one factor (2 levels) and model (4 levels) as the other, as described previously. These analyses tested the overall ability of these four proportional development models to predict later stage durations based on those of the earliest one. Also, since the ICD and EPD models were not found to best represent the majority of datasets tested (see Results, section 3) these tests allowed the ability of these simpler models to make predictions relative to the better but more complex ones to be tested, as well as whether the simpler models performed better on copepod data than on data for other arthropods.

## 3. Results

### 3.1. Distribution of development types among arthropods

Of the 71 arthropod species’ datasets examined, the development of only 9 (12.7 %) were best represented by an EPD model (Fig. 2). These included one arachnid [*Steneotarsonemus pallidus* (BANKS, 1901) data from Easterbrook et al. (2003)], five insects [*Anthonomus rubi* HERBST, 1795 and *Lygus rugulipennis* POPPIUS, 1911 data from Easterbrook et al. (2003), *Pseudococcus cryptus* HEMPEL, 1918 data from Kim et al. (2008), and *Frankliniella occidentalis* (PERGANDE, 1895) data from Nondillo et al. (2008)], and four crustaceans, all of which were calanoid copepods [*Eurytemora affinis*, *Pseudocalanus minutus*, and *Temora longicornis* data from Corkett and McLaren (1970) and *Pseudocalanus elongatus* (BOECK, 1865) data from Klein Breteler et al. (1995)]. Only one species dataset (1.4 %) was best represented by an ICD model of development (Fig. 2), specifically the dataset from Ouellet and Chabot (2005) for the northern shrimp, *Pandalus borealis*, a decapod crustacean.

**Figure 2.**
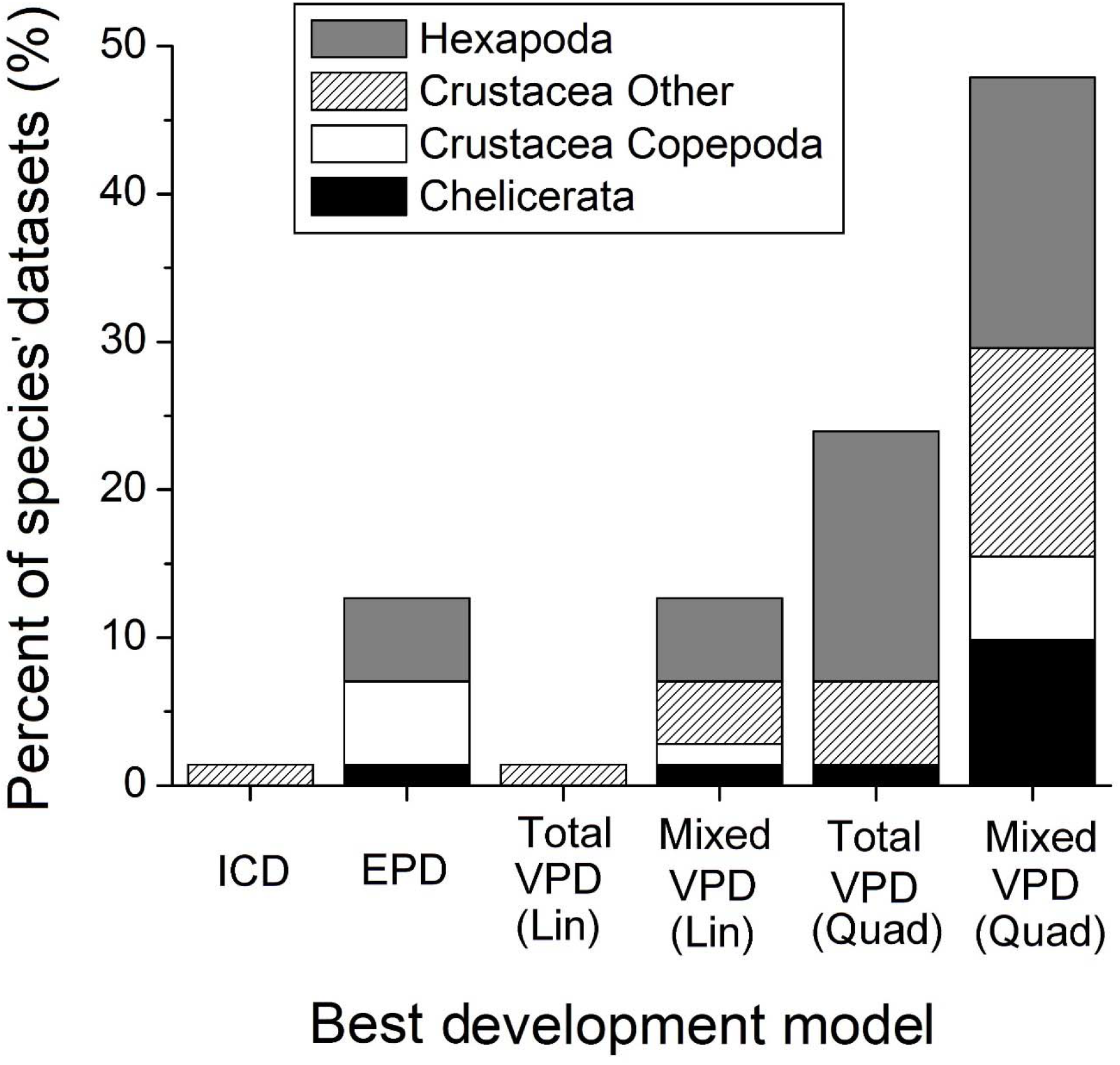
The percentage (%) of all species’ datasets (total = 71) for which each model of development was found to be the best (based on AIC_C_ ranking) in different arthropod groups. ICD = isochronal development; EPD = equiproportional development; VPD = variable proportional development, which could take either a linear (Lin) or quadratic (Quad) form, and be ‘ total’ (all stages’ proportions significantly affected by temperature) or ‘mixed’ (one or more stages’ proportions significantly affected by temperature, but not all stages).

Development of the remaining 61 species (see Supplementary File 3, Table S3) was best described by a ‘VPD’ model of development of either a linear (10 species = 14.1 % of datasets) or nonlinear quadratic (51 species = 71.8 % of datasets) form (Fig. 2). Of these 61 species for which either ‘VPD’ model was best, 18 represented cases of ‘total VPD’, in which the proportion of total development time taken up by all developmental stages varied significantly with temperature (Fig. 2). The remaining 43 species showed a ‘mixed’ form of development, in which the proportion of development taken up by one or more developmental stages significantly varied with temperature (representing stages with VPD), but proportions for one or more additional stages did not differ as a result of temperature (representing stages with EPD) (Fig. 2). Overall 5 crustaceans (all non-copepods), 1 arachnid, and 12 insects had total VPD, while 18 crustaceans (including 5 copepods), 8 arachnids, and 17 insects had mixed development.

Among species with mixed development, there was a tendency for more stages with relatively low numbers in the developmental sequence (1-4) to have their proportions significantly affected by temperature (Fig. 3A), but this was due to there being few species among those examined with more than 4 stages in their immature development (see Supplementary Files 2 and 3, Tables S3 and S4). When stage number was divided by the total number of stages for each species, this pattern disappeared and there were no obvious points in development when significant temperature effects were more prevalent (Fig. 3B). A possible exception to this result was the last developmental stage, for which 25 of the 43 species with mixed development (58.1 %) showed temperature effects on developmental proportions (Fig. 3B; see also Supplementary File 3, Table S4). However, the likelihood of the developmental proportion for a stage in a species with mixed development being affected by temperature was not significantly related to stage number (logistic regression pseudo-R^2^ = 0.001, stage number p = 0.606; Fig. 3A) or the proportion of development elapsed by the end of the stage (logistic regression pseudo-R^2^ = 0.001, development at stage p = 0.614; Fig. 3B). There were also no apparent differences among arthropod groups (Fig. 3A, B).

**Figure 3.**
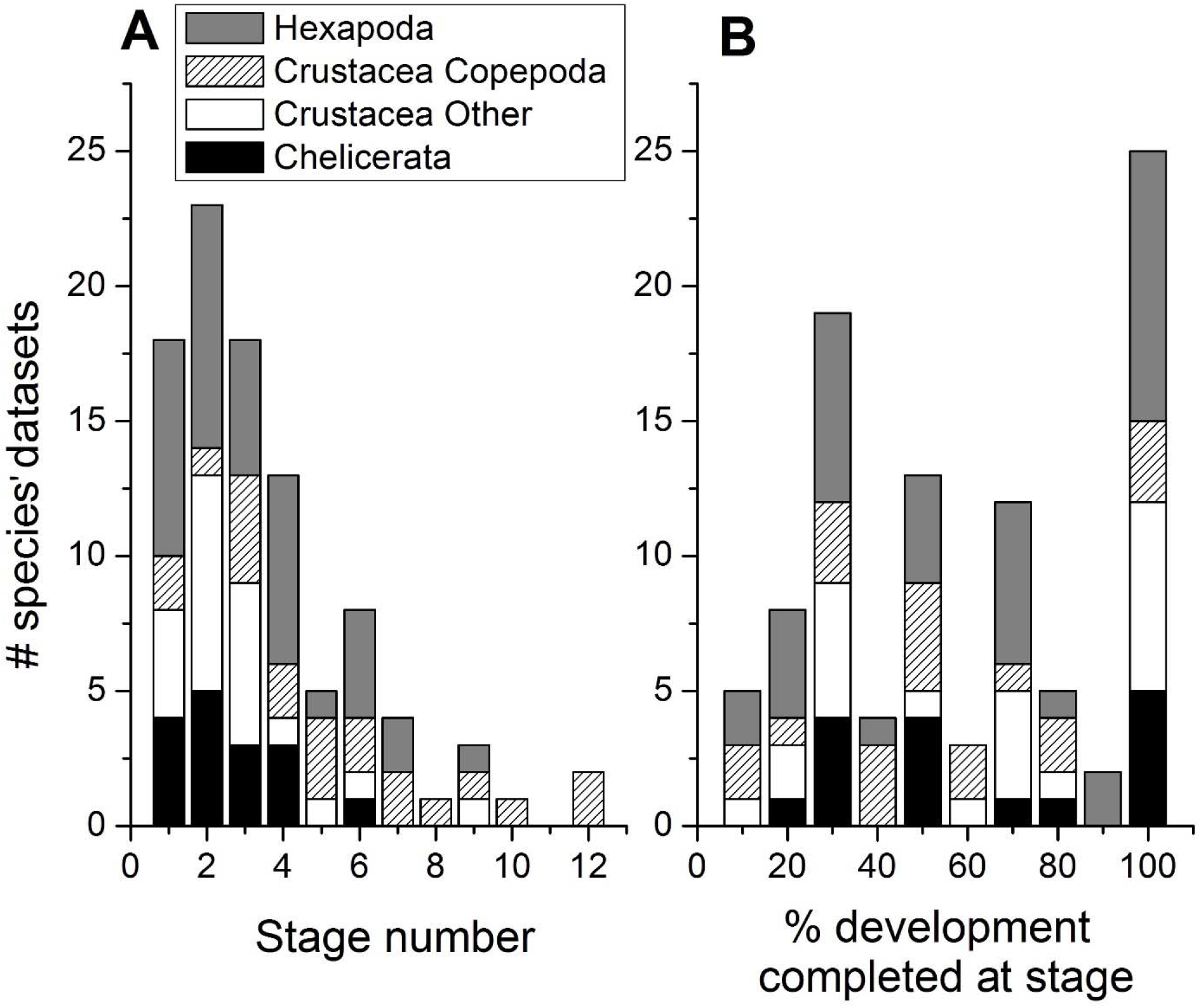
The number of species’ datasets with a mixed VPD type of development in which the developmental proportions for stages with different numbers (A, stages numbered from earliest to latest) or at which different percentages (%) of development were completed (B, calculated by dividing the stage number in A by the total number of stages in a species) were significantly affected by temperature. Data are divided up according to different arthropod groups.

### 3.2. Performance of different models of arthropod development

#### 3.2.1. Overall model performance

There were significant differences among the four models of arthropod development tested in terms of their ranking (out of four) based on AIC_C_-values (F _3, 272_ = 144.392, p < 0.001; Fig. 4A), model Δ_i_ (F _3, 272_ = 11.499, p < 0.001; Fig. 4B), pseudo-R^2^ values (F _3, 272_ = 142.392, p < 0.001; Fig. 4C), and cross-validation error (F _3, 272_ = 23.102, p < 0.001; Fig. 4D). There were no significant differences among arthropod subphyla (arachnids, crustaceans, and insects) in the performance of any of these functions assessed based on model ranking (Subphylum F _2, 272_ < 0.001, p > 0.999, Subphylum × Model interaction F _6, 272_ = 0.848, p = 0.534), Δ_i_ (Subphylum F _2,_ 272 = 2.481, p = 0.086, Subphylum × Model interaction F _6, 272_ = 1.705, p = 0.120), or pseudo-R^2^ values (Subphylum F _2, 272_ = 2.334, p = 0.099, Subphylum × Model interaction F _6, 272_ = 0.339, p = 0.916) (Fig. 4A-C). Cross-validation error did significantly vary among subphyla (F _2, 272_ = 10.835, p < 0.001), although there was no significant interaction between subphylum and model (F _6, 272_ = 0.845, p = 0.536) (Fig. 4D). This difference resulted because the cross-validation error of all models for crustaceans was significantly lower than that for insects (Tukey’s HSD test, p < 0.001; Fig. 4D), likely due to the better performance of most models on data for copepods (which are crustaceans) than non-copepods (see 3.2.2, Fig. 5). However, cross-validation error for arachnids did not significantly differ from that for insects (Tukey’s HSD test, p = 0.051) or crustaceans (Tukey’s HSD test, p = 0.650) (Fig. 4D).

**Figure 4.**
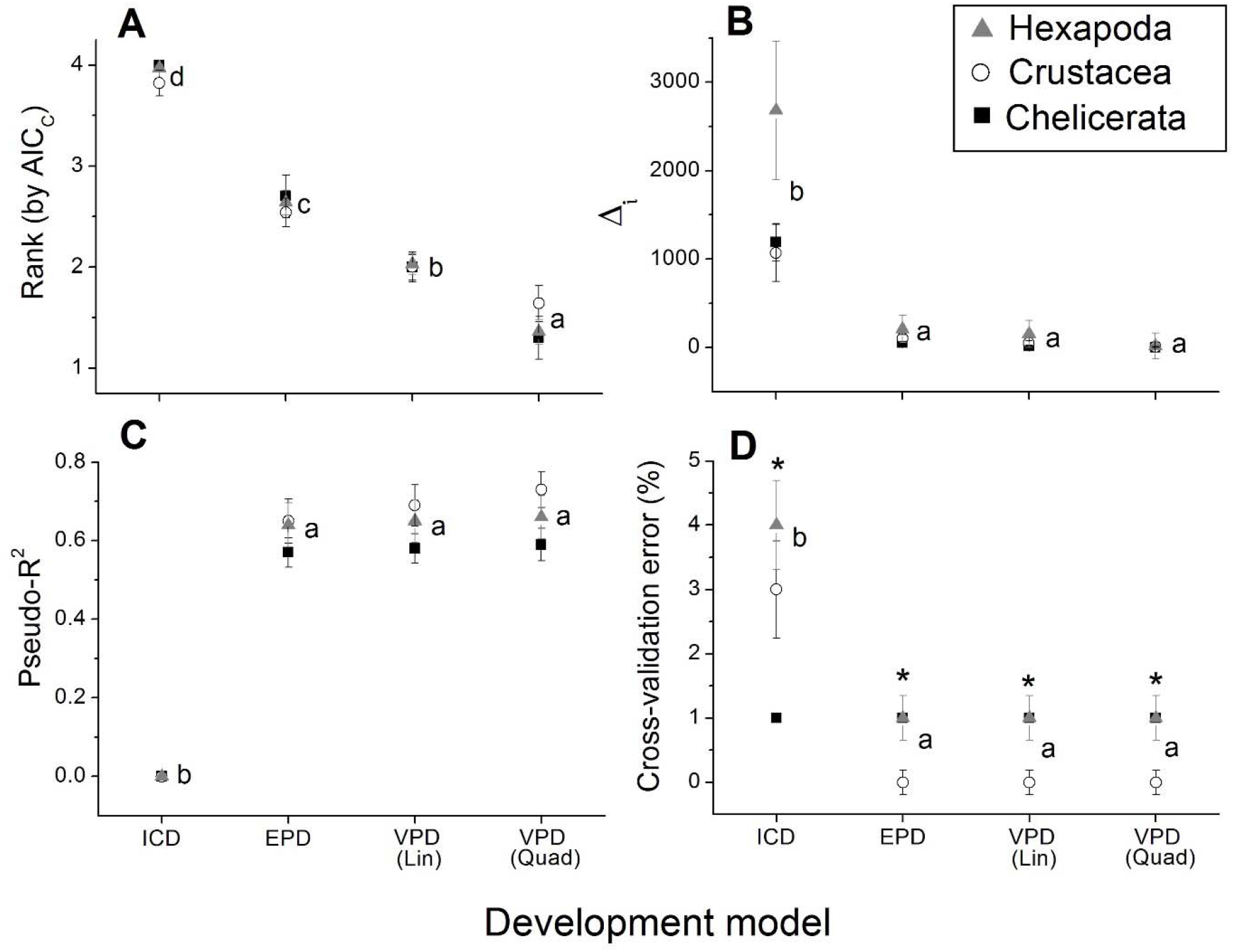
Performance of the different development models tested in this study (isochronal (ICD), equiproportional (EPD), and linear (Lin) and quadratic (Quad) forms of variable proportional (VPD) development), assessed based on each model’s (A) ranking based on AIC_C_ (1 = best, 4 = worst), (B) model Δ_i_ based on AIC_C_ values, (C) pseudo-R^2^, and (D) cross-validation error (%) when tested on data for each arthropod species. Mean ± SE values of each performance metric are plotted for each model and arthropod subphylum (chelicerates = black squares, crustaceans = white circles, and hexapods = gray triangles). Different lowercase letters to the right of the cluster of values for each model indicate models whose performance metrics varied significantly (Tukey’s HSD test, p ≤ 0.05), while an asterisk (‘*’) above these points indicates that performance differed significantly among subphyla (ANOVA, p ≤ 0.05).

**Figure 5.**
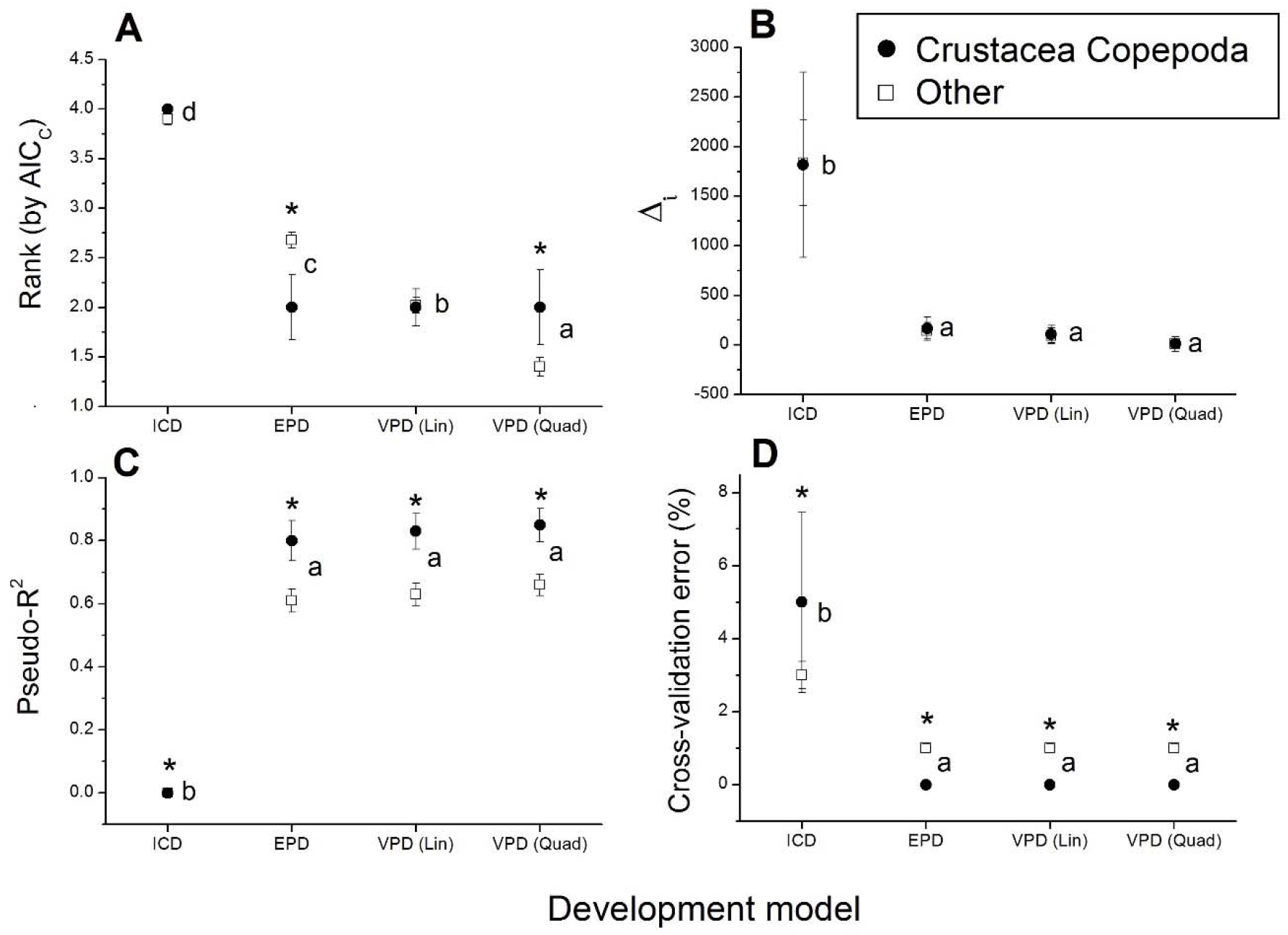
Performance of the different development models tested in this study (isochronal (ICD), equiproportional (EPD), and linear (Lin) and quadratic (Quad) forms of variable proportional (VPD) development), assessed based on each model’s (A) ranking based on AIC_C_ (1 = best, 4 = worst), (B) model Δ_i_ based on AIC_C_ values, (C) pseudo-R^2^, and (D) cross-validation error (%) when tested on data for copepod crustacean species (black circles) compared with those of all other arthropod taxa (white squares) species. Mean ± SE values of each performance metric are plotted for each model and group. Different lowercase letters to the right of the cluster of values for each model indicate models whose performance metrics varied significantly (Tukey’s HSD test, p ≤ 0.05), while an asterisk (‘*’) above these points indicates that performance differed significantly between copepod and non-copepod species (ANOVA, p ≤ 0.05).

Ranks of all models differed significantly (Tukey’s HSD test, all p < 0.001), with the lowest (best) rank overall achieved by the nonlinear quadratic form of the variable proportional (VPD) model, followed by the linear form of the VPD model, then the equiproportional model of development (EPD), and then lastly the isochronal model of development (ICD) (Fig. 4A). The ICD model was consistently the worst and drove the significance of differences among models, as it had significantly higher (meaning poorer) ranks (Fig. 4A), Δ_i_ (Fig. 2B), and cross-validation errors (Fig. 4D), and lower pseudo-R^2^ (Fig. 4C), than all other models (Tukey’s HSD test, all p < 0.001). Conversely, Δi, pseudo-R^2^, and cross-validation error did not significantly differ among EPD and both forms of the VPD models (Tukey’s HSD test, p ≥ 0.598; Fig. 4), and removing the ICD model from analyses resulted in non-significant differences among models in terms of all metrics except rank (results not shown).

#### 3.2.2.

Comparison of performance between copepod and non-copepod species’ data

Differences in the performance of models assessed in terms of Δ_i_, (Model: F _3, 276_ = 7.034, p < 0.001; Model × Copepod: F _3, 276_ = 0.001, p > 0.999; Fig. 5B), pseudo-R^2^ values (Model: F _3,_ 276 = 98.453, p < 0.001; Model × Copepod: F _3, 276_ = 1.295, p = 0.276; Fig. 5C), and cross-validation error (Model: F _3, 276_ = 24.603, p < 0.001; Model × Copepod: F _3, 276_ = 2.069, p = 0.105; Fig. 5D) were consistent between copepod and non-copepod arthropod species. Model cross-validation errors were significantly lower (F _1, 276_ = 4.089, p = 0.044; Fig. 5D) and pseudo-R^2^ values were significantly higher (F _1, 276_ = 11.650, p = 0.001; Fig. 5C) for copepods than non-copepod species. However, model Δ_i_ did not differ significantly between copepods and non-copepods (F _1, 276_ < 0.001, p = 0.988; Fig. 5B). There was a significant interaction between the effects of model type and whether species were or were not copepods on model rankings (F _3, 276_ = 4.843, p = 0.003; Fig. 5A). This interaction resulted because certain models were ranked better overall for copepod species data than they were for non-copepods, specifically the quadratic VPD (one-way ANOVA comparing copepods versus non-copepods for one model at a time: F _1,_ 69 = 4.137, p = 0.046) and EPD models (F _1, 69_ = 7.209, p = 0.009; Fig. 5A), while rankings for the linear VPD (F _1, 69_ = 0.005, p = 0.944) and ICD models (F _1, 69_ = 0.331, p = 0.567) did not significantly differ between copepod and non-copepod species (Fig. 5A).

### 3.3. Prediction error of development models

Absolute percent prediction error differed significantly among models (F _3, 272_ = 31.144, p < 0.001), but not among subphyla (Subphylum: F _2, 272_ = 0.237, p = 0.789; Model × Subphylum: F _6,_ 272 = 1.739, p = 0.112) (Fig. 6A). Differences among models were driven by the fact that ICD (mean percent error = 77.8 %) had much and significantly (Tukey’s HSD test, all p < 0.001) greater error than all other models (mean errors = 25.0-28.4 %) (Fig. 6A). EPD and both forms of VPD did not have significantly different percent errors from each other (Tukey’s HSD test, p ≥ 0.948) (Fig. 6A). If ICD was removed from consideration, there was no significant difference among models in terms of their prediction error (e.g., Model F _2, 204_ = 0.250, p = 0.772; other results not shown). The prediction errors of the ICD, EPD, linear VPD, and quadratic VPD models were overall not significantly greater than 64, 23, 21, and 19 %, respectively (one-sample *t*-tests, t _70_ ≤ 1.921, p ≥ 0.056). When comparing model predictive ability on copepod versus non-copepod species’ datasets, percent errors also did not differ between copepods and non-copepod species (Copepod: F _1, 276_ = 1.212, p = 0.272; Model × Copepod: F _3, 276_ = 0.141, p = 0.936), only among models (F _3, 276_ = 16.558, p < 0.001) (Fig. 6B).

**Figure 6.**
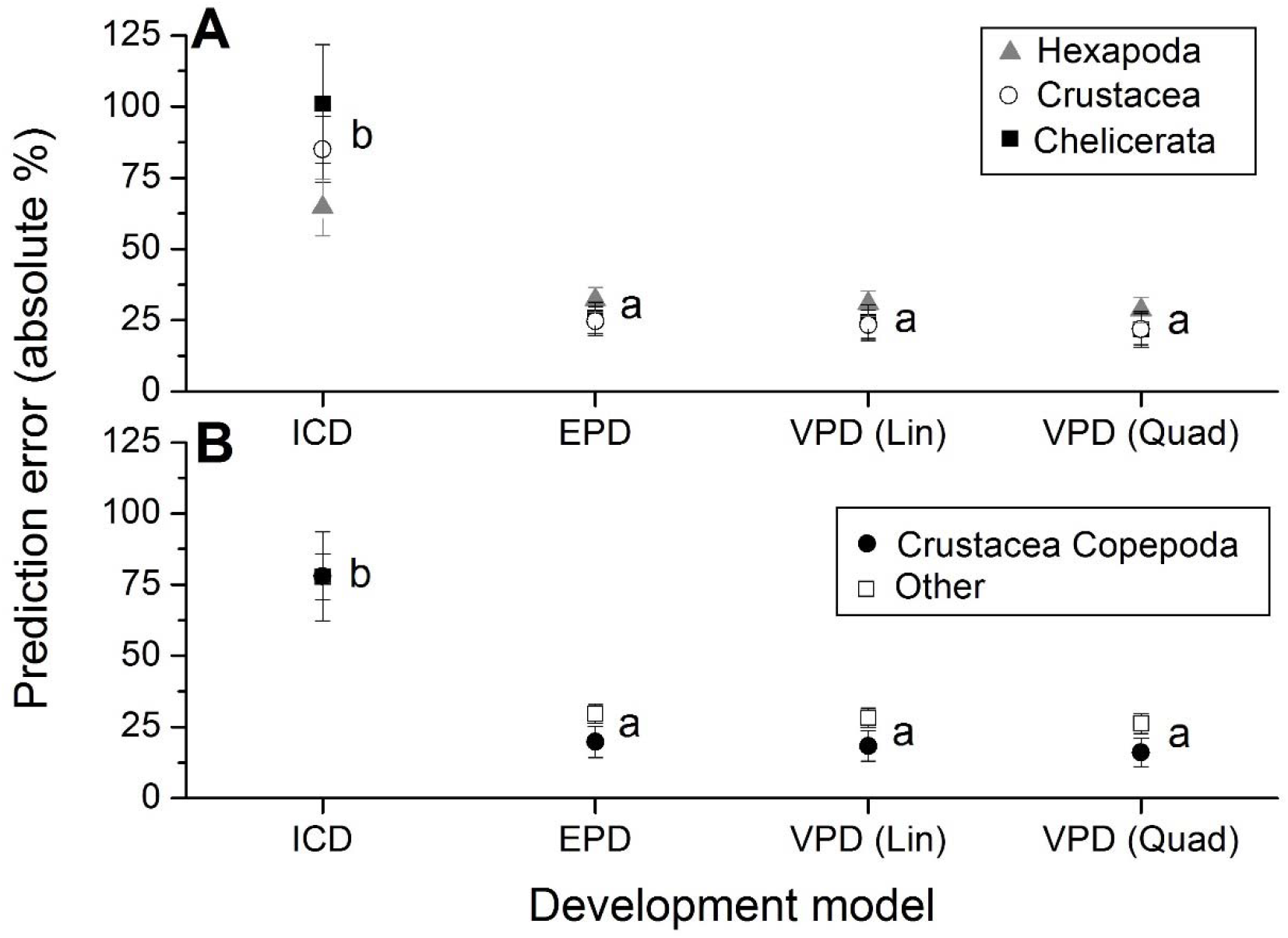
Prediction errors of the different development models tested in this study (isochronal (ICD), equiproportional (EPD), and linear (Lin) and quadratic (Quad) forms of variable proportional (VPD) development). An average absolute percent (%) error was calculated for each species’ dataset by comparing durations predicted based on each model for all later stages based on the duration of the first stage (see section 2.5 for details) to the actual observed durations for these stages in original studies. Mean ± SE values of prediction error calculated across species are plotted for each model and (A) arthropod subphylum (chelicerates = black squares, crustaceans = white circles, and hexapods = gray triangles) or (B) for copepod crustaceans (black circles) and all other arthropod taxa (white squares). Different lowercase letters to the right of the cluster of values for each model indicate models whose prediction errors differed significantly (Tukey’s HSD test, p ≤ 0.05); prediction errors did not differ significantly among subphyla or between copepod and non-copepod species (ANOVA, p > 0.05).

## 4. Discussion

### 4.1. Comparison of results to previous studies

In the present study, it was found that isochronal (ICD) and equiproportional development (EPD) were not general phenomena among the arthropod species examined, as the development of the vast majority of species (61 out of 71, or 85.9 %) was best represented by a variable proportional (VPD) model of development. In comparison with previous studies of some of the species examined in this one, the new analytical technique used led to different conclusions regarding whether they had ICD or EPD. The decapod crustaceans *Chionoecetes opilio*, *Pandalus borealis*, and *Pandalus montagui* were previously suspected to have EPD (*C. opilio* and *P. borealis*: P. Ouellet and B. Sainte-Marie, Fisheries and Oceans Canada, pers. comm.) or ICD (*P. montagui*:Miller, 2008), but in this study *C. opilio* and *P. montagui* were concluded to have VPD, and *P. borealis* was the only species concluded to have ICD. Both MacKenzie (1988) and Quinn et al. (2013) concluded that the decapod *Homarus americanus* had EPD, but reanalyses of their data in the present study showed that this species has VPD, with the proportions of development in all three zoeae and the decapodid stage (MacKenzie, 1988) or the third zoea only (Quinn et al. 2013) varying with temperature in a quadratic manner (Supplementary File 3, Tables S3 and S4). Results of this study did, however, confirm those inferred from the different developmental proportions reported at different temperatures by Lefebvre and Pasquerault (2004) for the insect *Ophyra aenescens*, as this species was found to have VPD. Among the 9 copepods examined in the present study, the development of four species (*Eurytemora affinis*, *Pseudocalanus elongatus* (data from Klein Breteler et al., 1995)*, Pseudocalanus minutus*, and *Temora longicornis*) previously thought to have EPD (Corkett and McLaren, 1970; Hart, 1990; Kiørboe and Sabatini, 1994; Petersen, 2001) were concluded to be best represented as EPD. However, the remaining five copepod species’ datasets were all concluded to be best represented by the VPD models, and three of these species were concluded by previous studies (Miller et al., 1977; Geiling and Campbell, 1992; Hart, 1990; Kiørboe and Sabatini, 1994; Petersen, 2001) to have EPD [*Skistodiaptomus pallidus* (HERRICK, 1879) data from Kamps (1978), and *Pseudocalanus elongatus* data from Thompson (1982)] or ICD [*Acartia (Acanthacartia) tonsa* DANA, 1849 data from Leandro et al. (2006b)]) (see Supplementary File 1, Table S1). Therefore, while EPD did seem to represent data for copepod species somewhat better than that of other crustacean and arthropod taxa, the general occurrence of EPD among copepod crustaceans noted in previous studies was not confirmed.

The different conclusions from previous studies described above were presumably due to the use of an improved method to test for EPD in this study, as previous studies either did not perform formal statistical tests for ICD or EPD, or if they did, they mostly did not also test and compare alternative models (e.g., VPD). The differences in conclusions for some datasets for the same species but obtained from different studies [*H. americanus*: total VPD (MacKenzie, 1988) vs. mixed VPD (Quinn et al., 2013); *P. elongatus*: EPD (Klein Breteler et al., 1995) vs. VPD (Thompson, 1982)] were interesting outcomes. It is noteworthy that in each of these pairs of studies, the one for which the least variable form of development was concluded as best was also the one with the smaller sample size of the two (Supplementary File 2). The ten species for which ICD or EPD were concluded to be a better representation of their development than VPD also tended to have relatively small sample sizes for each stage and temperature compared to all the other datasets examined (see Supplementary File 2). Therefore, it is very likely that the conclusion of EPD in these cases does not represent an actual biological characteristic of these species, but simply insufficient power to detect their VPD due to low replication. Indeed, if sample sizes are small but the models fit to the data become more complex, as in the case of VPD (especially the quadratic form of the model), the bias introduced by greater model complexity (i.e. more parameters) outweighs the improved fit achieved by the model, and thus lowers its information content, increases its AIC_C_ value, and worsens its rank (Anderson, 2008; Quinn, 2017). It would thus be reasonable to expect an analysis of data from a better-replicated study of the species for which ICD and EPD were concluded in this study to result in them being found to actually have VPD.

### 4.2. The reality of ICD and EPD and implications of plastic and mixed development

The occurrence of ICD and EPD in the immature development of arthropods, including among the copepod species for which they were first reported, is thus unlikely to be a ‘real’ phenomenon in nature. This finding is perhaps not so surprising, given that an increasing body of work in recent years has shown that many (and possibly all?) arthropod species exhibit at least some plasticity in their immature developmental pathways (Jarošík et al., 2002, 2004; Oliphant et al., 2013; Karimi-Malati et al., 2014; Quinn, 2016). Many species of decapod crustaceans (e.g., Boyd and Johnson, 1963; Hamasaki et al., 2009) and insects (e.g., Karimi-Malati et al., 2014) develop through additional or alternative stages or instars under suboptimal conditions, such as extreme temperatures, low food supply, presence of contaminants, or limited availability of substrate for settlement (Oliphant et al., 2013; Quinn, 2016 and references therein). In many cases, such developmental plasticity is thought to be adaptive; for example, at low temperatures the shrimp *Palaemon varians* LEACH, 1813 (formerly *Palaemonetes varians*) develops through 1-3 additional stages at lower temperatures than it does at higher temperatures, which is presumed to allow it to delay settlement to juvenile habitats and give it more time to feed and grow to a larger size before settling (resulting in lower post-settlement mortality) than it otherwise could achieve under low-temperature conditions (Oliphant et al., 2013). In many of the species examined herein and found to have VPD, additional developmental stages did occur at some temperatures (see original studies cited in Supplementary File 1, Tables S2), resulting in immediately altered proportions for all ‘normal’ stages at these temperatures. However, for many others with VPD extra stages did not occur, but instead the specific allocation of the developmental period to each stage changed among temperatures (e.g., Campolo et al., 2004). The specific implications of this form of plasticity are not immediately clear and likely vary on a case-by-case basis, but one can imagine similar outcomes to the occurrence of extra stages. For instance, the lengthening of the last developmental stage at suboptimal temperatures may permit more feeding in preparation for metamorphosis, and thus attaining a larger postmetamorphic size (*sensu*Oliphant et al., 2013). The results of the present study strongly suggest that developmental plasticity is a general phenomenon among arthropods, but more research is needed on the exact nature, evolution, and implications of developmental plasticity among arthropod taxa (Quinn, 2016).

The observation of a ‘mixed’ type of development between EPD and VPD in many species in the present study was an unexpected finding, which suggests that even within the same species the proportion of development spent in different stages may be differently affected (or not) by temperature. No obvious overall relationship between where in the developmental sequence a stage fell and the likelihood of its proportion being affected by temperature was found, so the reasons for the temperature (in)dependence of each species’ stages’ developmental proportions must be investigated individually in a future analysis. One can imagine that some stages likely have greater scope for developmental plasticity than others. For example in a species with both feeding and non-feeding stages, one would expect the non-feeding stages to have less plasticity due to their lifetime being limited by stored reserves (Jirkowski et al., 2015; Pineda and Reyns, 2018). Perhaps some intermediately timed stages might also be less flexible if processes occur during them that are required for later development and successful metamorphosis, as in the second zoea stage of many decapods (Anger 2001; Quinn, 2016). Later stages of crustaceans, which must settle to the benthos before metamorphosing, may also be more plastic due to the possibility of them not encountering suitable substrate immediately when reaching competence (Pineda and Reyns, 2018) or having experienced insufficient growth as larvae to be successful after metamorphosis (Oliphant et al., 2013). Future analyses of such possible features of mixed and VPD types of development will no doubt reveal interesting characteristics of arthropod species.

### 4.3. Possible predictive utility of EPD

Using the VPD models resulted in better overall predictive ability of later developmental stages’ durations than did EPD, and especially ICD, so it does represent an improvement over these simpler models that should be used more in future studies. However, an important finding of this study was that although EPD was rarely the best descriptor of any species’ immature development, it did perform comparably well to the VPD models on most species’ datasets and had similar predictive ability to these ‘better’ models. The mean pseudo-R^2^ values of the EPD model were always within 1-8 % of those of the VPD models (depending on subphylum), and the EPD model had equal mean cross-validation error to the VPD models. The mean prediction error of the EPD model was also within 1-4 % of that of the two VPD models, and all three of these models had overall prediction errors significantly less than or equal to 19-23 %. Therefore, although EPD is likely not ‘real’ and is not the best representation of most species’ immature development, for most practical purposes it may be appropriate to use it as a fair approximation thereof. This is useful, as obtaining sufficient data to model a species’ development as VPD would require a lot of time and resources, whereas the data required to model EPD and make predictions of later stages’ durations based on it is potentially much less (see Fig. 1B, F in section 1; Corkett, 1984; Hart, 1990). Caution should still be taken before applying EPD too broadly, however, as the actual nature of arthropod development is plastic (VPD), and in some cases (e.g., occurrence of extra larval stages and/or plasticity near extreme temperatures; Oliphant et al., 2013; Karimi-Malati et al., 2014) the developmental alterations that result from this plasticity can have major impacts on the ecology of immature stages (Oliphant et al., 2013; Quinn, 2016). Therefore, the decision of whether predictions made using EPD are accurate enough must lie with each individual study, and be based on its objectives and the sensitivity of these outcomes to the developmental plasticity of the species involved.

### 4.4. Limitations and perspectives for future work

There are of course some important limitations to the results of the present study. While the number of species’ datasets examined (71) was fairly large and included members of all three major arthropod subphyla, these only provided information for species from a limited sample of the diverse taxa within these subphyla, including only two crustacean classes, very few copepod species, and only 10 chelicerates (all arachnids, all but one of which were mites). These datasets were also obtained in a non-random and semi-arbitrary manner from studies examined in previous reviews (Quinn, 2016, 2017), although it should be noted that Quinn (2017) did perform a standardized search of the temperature-dependent arthropod development literature to obtain their study datasets. For a small percentage of datasets (14.1 %), total developmental durations were not reported in original studies and thus had to be indirectly calculated, which may have introduced some errors (though probably small ones) into the calculation of developmental proportions. While a more extensive survey of the literature encompassing more arthropod species and taxa would certainly provide stronger conclusions (and should certainly be done, see below), there was no reason to suspect any particular biases in the results obtained herein affecting the main conclusions. Further, the fact that similar trends, such as VPD being far more prevalent than ICD or EPD, were found for members of all three subphyla examined lends some support to the generality and representativeness of the conclusions drawn.

It is also important to note that the results for copepods herein strongly call in to question the occurrence of ICD and EPD among these crustaceans, which is contrary to a great deal of previous research (e.g., Hart, 1990, 1994, 1998; Petersen, 2001; but see Carlotti and Nival, 1991; Petersen and Painting, 1990). However, to confirm this result more than nine species of copepods need to be examined. If possible, a follow-up review should attempt to obtain more high-quality data from the literature for many more copepod species, including for more of those that were previously concluded to have ICD or EPD, and test their data using the techniques developed in this study. Given the types of data (e.g., means only) based on which previous studies concluded ICD and EPD in copepods, however, extracting sufficient data from published papers to test with VPD models may not be possible; in this case, new experimental studies rearing copepods at different temperatures will be necessary to obtain the data needed to conclusively test whether ICD or EPD rather than VPD occur among these species.

Perhaps a more significant limitation was that a wide variety of developmental increments were equated under the common term ‘developmental stages’ in the present study (Table 1). There are important distinctions between developmental phases, stages, and instars, for example, which were ignored in this study, but as outlined earlier (see section 2.1) it is unlikely that these inconsistencies should have affected the conclusions of the present study. However, a study accounting for phase-stage-instar differences in developmental proportions may be illuminating, and thus should be done in the future. The possibility that the type of development or scope for developmental plasticity might differ among arthropods with anamorphic as opposed to metamorphic development may also be worth investigating.

The statistical analyses performed in the present study were robust and novel, but they did also have limitations that could be improved in future work. For example, the linear and quadratic VPD models used are only two of a large number of different potential temperature-dependent development function types (e.g., Quinn, 2017) that might be applied to such data, and some more complex function types exist that can even capture underlying physiological processes, such as enzyme thermoactivation and thermoinhibition, involved in thermal effects on development (e.g., Schoolfield et al., 1981; Shi et al., 2017c). While such highly complex functions could only be applied to datasets with relatively large sample sizes, it would certainly be worth looking at developmental proportions using them in the future. It is also conceivable that the developmental proportions of stages in some species may not exhibit an even, curvilinear relationship with temperature, but might instead vary in a ‘chaotic’ or ‘sporadic’ manner among temperatures. A reasonable model of ‘sporadic’ development could not be developed in the present study, but it may be worth pursuing in the future. It is also well established that development times of arthropods vary not only with temperature, but also as a function of other environmental and biotic factors such as salinity, food quality or quantity, social interactions, oxygen, pH, etc., and different combinations thereof (Anger, 1991; Klein Breteler, 1980; Klein Breteler et al., 1995; Anger, 2001; Keppel et al., 2012); the possibility that developmental proportions might vary due to such factors should also be investigated using an approach analogous to the one developed for temperature in this study. It would also likely be very useful to combined the datasets and approaches used in this study with some of those from the literature on development rate isomorphy among ectotherms (Jarošík, 2002, 2004; Shi et al., 2010; Sandhu et al., 2011; Kuang et al., 2012; Lin et al., 2018). Doing so could increase the taxonomic range over which data were collected and analyzed, and allow for these two approaches to be combined and analyzed in interesting and useful ways.

Analyses in this study were also confined to the available data obtained from previous studies, and thus all assessments of model performance and predictive ability were based on the fit of development models to the data used to generate them. Therefore, a true test of the performance and predictive ability of models of different types of development would need to be carried out on new data (Picard and Cook, 1984). For example, future studies should rear immature stages of the species examined in this one, and compare the developmental proportions and durations of different stages at different temperatures to the stage- and temperature-specific developmental proportions for these species predicted by the different models (EPD, VPD, etc.) tested in this study. Only then could a definitive conclusion be reached regarding the occurrence or not of EPD and other development types among arthropods be reached. Of course, such studies should also test other factors impacting development times (salinity, etc. as described above), and likely also should be conducted over very wide thermal ranges to determine whether and how developmental proportionality breaks down near thermal extremes (Schoolfield et al., 1981; Shi et al., 2012; Quinn, 2017). Also, since such studies are most likely to be conducted in laboratory settings, at constant temperatures, caution should be taken in applying their results (and those of the present study) to predicting arthropod development in nature. Development in variable thermal regimes can differ from that at constant temperatures (Anger, 1983), and in field conditions development can occur more quickly or slowly (Anger, 2001; Niehaus et al., 2012) than in the laboratory, depending on the species.

A considerable amount of information on the proportions of development spent by arthropod species belonging to various taxa in different immature stages at different temperatures has been amassed during the present study (Supplementary File 3, Table S4). As already discussed, these data and the developmental proportions predicted by models of ICD, EPD, and VPD based on them could be used to predict development times in future studies on these species. These data also provide information useful for endeavours such as larval dispersal modeling (Reitzel et al., 2004; Quinn et al., 2013; Pineda and Reyns, 2018), predictions of pest dynamics (Easterbrook et al., 2003; Campolo et al., 2009; Eliopoulous et al., 2010), ecosystem models (Huntley and López, 1992), and so on. However, there is also much potential for this information on the developmental proportions and their dependence on temperature (or not) to be analyzed across species and higher taxa from an ecological or evolutionary perspective.

The proportion of development spent by a particular species in each developmental stage may reflect evolutionary adaptation to the phenology of conditions encountered in its habitat. For example, most decapod crustacean larval stages take up progressively longer portions of development (MacKenzie, 1988; Anger, 2001; see also the data in Supplementary Files 2 and 3). In decapod species in temperate regions, this means that early-stage larvae, which take up the smallest proportion of development at a particular temperature, experience the coldest conditions, while later-stage larvae that can take up larger developmental proportions experience the warmest conditions; this means that the shortest, earliest stages experience the greatest inhibition by cold, while the longest, later stages experience the greatest acceleration by warmth, which may optimize overall development time and effectively make development in nature isochronal. As reported for copepods (Hart, 1994), the proportions for a particular species’ stages might also be optimized to avoid temporal and spatial overlap with competing species in the same habitat. It is also conceivable that the degree of developmental plasticity exhibited by a species can be explained by its life habits. Species inhabiting environments with more variable temperature regimes than others, such as terrestrial species, might be expected to have more flexible development than those in less variable (e.g., aquatic) habitats (Shi et al., 2012; Quinn, 2017). There is thus the potential to compare the characteristics of one species’ development with its habitat to assess whether particular features of the habitat have selected for a particular form of development.

While no obvious taxonomic patterns were detected in the present study, the information collected might later be used on a larger scale to specifically answer broad evolutionary questions. Heterochrony is the changing of the timing of events during development, and it is associated with many major evolutionary transitions, including among arthropods (Jirkowsky et al., 2015). A phylogenetic analysis of the proportion of development spent in different stages among closely related species or higher taxa might reveal interesting evolutionary patterns. For example, one could examine whether: the loss or gain of a larval stage is associated with changes to the developmental proportions of other stages; greater plasticity is associated with species shifting their ranges into more thermally variable environments; developmental proportions can be used in addition to other phylogenetic characters to enhance taxonomical distinctions among species; evolution towards direct development (loss of free-living larval stages) is preceded by reductions in developmental proportions of larval stages; apparently similar larval stages, such as the nauplii of dendrobranchiate and euphausiid crustaceans, are equivalent to those that are free-living in other taxa or embryonized (reduced and completed in the egg) in others (Jirkowski et al., 2015); and so on. While a more extensive database including many more species and higher taxa is needed before such analyses can be conducted, the present study is a first step in this direction, which should lead to many informative and interesting investigations in the future.

## Acknowledgements

The University of New Brunswick, Saint John Campus, provided resources that made the review and analyses in this paper possible. I thank Gerhard Scholtz and two anonymous reviewers for comments that improved the manuscript. The author thanks Bernard Sainte-Marie, Patrick Ouellet, Heather Major, Heather Hunt, Jeff Houlahan, Rémy Rochette, and members of the Biology 6000 class feedback committee and Rochette-Hunt lab at UNBSJ for feedback and guidance during this project. I thank Tammy Sha Bo in particular for observations that led to the idea and models for VPD developed herein, which allowed for much more sensible and novel tests of EPD versus alternatives; this work is dedicated to her.

## Electronic Supplementary Materials

**Supplementary File 1.** Contains Tables S1 and S2, along with their associated reference lists.

**Table S1.** Species of copepods (subphylum Crustacea: class Hexanauplia: subclass Copepoda: infraclass Neocopepoda) that previous studies concluded or suggested had either ICD or EPD. The information in this table was obtained through the literature searches (Web of Science and Google Scholar) discussed in the Introduction (section 1) of the main text.

**Table S2.** Arthropod species for which datasets were obtained from previously published studies and used in analyses within this one.

**Supplementary File 2.** Contains the datasets extracted for each species from each study (‘Input Data’), an R code (‘EPD tests code’) written by the author to test each dataset for ICD, EPD, and linear and quadratic forms of VPD and produce AIC_C_ values, cross-validation errors, and summary output for each of these development models, and the R output of analyses performed with this code on each species’ dataset (‘Output Data’).

**Supplementary File 3.** Contains Tables S3 and S4.

**Table S3.** Summary of performance metrics calculated for each development model [ICD, EPD, VPD (Lin), and VPD (Quad)] for each species’ dataset tested, including model AIC_C_, Δ_i_, rank based on AIC_C_ (1-4), cross-validation error, pseudo-R^2^, and mean raw (in d) and percent (%) absolute prediction errors calculated for each species’ dataset. Whether conditions for concluding ICD or EPD were met by the data is also outlined. Whether original studies reported full development times and if each species had anamorphic or metamorphic development is also listed. The number of developmental stages whose proportions were significantly affected by temperature or not is also listed, as well as the proportion of stages with significant temperature effects, and thus whether or not the species had mixed development. The studies cited in this table are provided with Table S2 in Supplementary File 1.

**Table S4.** The proportions of total development spent by each of the species examined in this study in each of their immature developmental stages, as predicted based on the parameters of development models fit to them using GLMs in the present study. For all equations, the predicted proportions were back-transformed from arcsine-square-roots. Equations are included as Excel formulae, and those for VPD models are linked to values in the first column representing temperature (set to an arbitrary value of 15°C in this file). Whether the developmental proportion of each stage was significantly affected by temperature is also indicated (1 = affected, 0 = not affected) in the columns to the right of each VPD model’s column. Whether each developmental stage was feeding or non-feeding is also listed. The studies cited in this table are provided with Table S2 in Supplementary File 1.

